# Common maternal and fetal genetic variants show expected polygenic effects on the probability of being born small- or large-for-gestational-age (SGA or LGA), except in the smallest 3% of babies

**DOI:** 10.1101/2020.03.25.005660

**Authors:** Robin N Beaumont, Sarah J Kotecha, Andrew R. Wood, Bridget A. Knight, Sylvain Sebert, Mark I. McCarthy, Andrew T. Hattersley, Marjo-Riitta Järvelin, Nicholas J. Timpson, Rachel M Freathy, Sailesh Kotecha

**Affiliations:** Institute of Biomedical and Clinical Science, University of Exeter, Exeter, UK; Department of Child Health, School of Medicine, Cardiff University, Cardiff, UK; Center For Life-course Health Research, University of Oulu, Finland; Department of Epidemiology and Biostatistics, Imperial College, London, UK; Wellcome Trust Centre for Human Genetics, University of Oxford, Oxford, OX3 7BN, UK; Oxford Centre for Diabetes, Endocrinology and Metabolism, University of Oxford, Oxford, OX3 7LE, UK; Oxford National Institute for Health Research (NIHR) Biomedical Research Centre, Churchill Hospital, Oxford, OX3 7LE, UK; Medical Research Council Integrative Epidemiology Unit, University of Bristol, Bristol, UK

## Abstract

Babies born clinically Small- or Large-for-Gestational-Age (SGA or LGA; sex- and gestational age-adjusted birth weight (BW) <10^th^ or >90^th^ percentile, respectively), are at higher risks of complications. SGA and LGA include babies who have experienced growth-restriction or overgrowth, respectively, and babies who are naturally small or large. However, the relative proportions within each group are unclear. We aimed to assess the extent to which the genetics of normal variation in birth weight influence the probability of SGA/LGA. We calculated independent fetal and maternal genetic scores (GS) for BW in 12,125 babies and 5,187 mothers. These scores capture the direct fetal and indirect maternal (via intrauterine environment) genetic contributions to BW, respectively. We also calculated maternal fasting glucose (FG) and systolic blood pressure (SBP) GS. We tested associations between each GS and probability of SGA or LGA. For the BW GS, we used simulations to assess evidence of deviation from an expected polygenic model.

Higher BW GS were strongly associated with lower odds of SGA and higher odds of LGA (OR_fetal_=0.65 (0.60,0.71) and 1.47 (1.36,1.59); OR_maternal_=0.80 (0.76,0.87) and 1.23 (1.15,1.31), respectively per 1 decile higher GS). Associations were in accordance with a polygenic model except in the smallest 3% of babies (P_fetal_=0.0034, P_maternal_=0.023). Higher maternal GS for FG and SBP were associated with higher odds of LGA and SGA respectively (both P<0.01). While lower maternal FG and SBP are generally considered healthy in pregnancy, we found some evidence of association with higher odds of SGA (P=0.015) and LGA (P=0.14) respectively.

We conclude that common genetic variants contribute to risk of SGA and LGA, but that additional factors become more important for risk of SGA in the smallest 3% of babies. Naturally low maternal glucose and blood pressure levels may additionally contribute to risk of SGA and LGA, respectively.

**Author Summary:** Babies in the lowest or highest 10% of the population distribution of birth weight (BW) for a given gestational age are referred to as Small- or Large-for-Gestational-Age (SGA or LGA) respectively. These babies have higher risks of complications compared to babies with BW closer to the mean. SGA and LGA babies may have experienced growth restriction or overgrowth, respectively, but may alternatively just be at the tail ends of the normal growth distribution. The relative proportions of normal vs. sub-optimal growth within these groups is unclear. To examine the role of common genetic variation in SGA and LGA, we tested their associations with a fetal genetic score (GS) for BW in 12,125 European-ancestry individuals. We also tested associations with maternal GS (5,187 mothers) for offspring BW, fasting glucose and systolic blood pressure, each of which influences fetal growth via the *in utero* environment. We found all fetal and maternal GS were associated with SGA and LGA, supporting strong maternal and fetal genetic contributions to birth weight in both tails of the distribution. However, within the smallest 3% of babies, the maternal and fetal GS for BW were higher than expected, suggesting factors additional to common genetic variation are more important in determining birth weight in these very small babies.

## Introduction

Size at birth is an important factor in new-born and infant survival. Term-born babies are most frequently admitted to the neonatal unit when born at the extremes of the birth weight distribution [1]. Small for Gestational Age (SGA; defined as birth weight adjusted for sex and gestational age that is below the 10^th^ percentile of the population or customized standard) is often used as a proxy indicator of fetal or intrauterine growth restriction (FGR or IUGR [2]). A fetus is described as growth-restricted when it has failed to reach its growth potential due to impaired placental function [3] or due to fetal or maternal reasons, and SGA fetuses are at higher risk of adverse outcomes such as stillbirth [4]. It is likely that SGA infants who are genetically small are at a lower risk of future morbidity than FGR infants. Risks of adverse outcomes are increased in preterm babies, and the mechanisms associated with SGA in preterm babies are likely to be different to those born at term. SGA and FGR are often used interchangeably since fetal growth can be difficult to measure. [2,3] However, they are not synonymous: not all growth-restricted fetuses are small enough to be considered SGA [5], and the SGA group itself is heterogeneous with an estimated 50-70% being constitutionally small babies with normal placental function and outcomes [6], in addition to babies expected to be small due to chromosomal anomalies [7].

At the upper end of the birth weight distribution, large for gestational age (LGA, defined as sex- and gestational age-adjusted birth weight >90^th^ percentile) is associated with a higher risk of obstructed labour, which can lead to complications for both mother and baby, including injury, neonatal hypoglycaemia and even fetal death [8]. LGA may indicate excessive growth of the fetus, for example due to elevated maternal glycemia, which is a major determinant of fetal growth [9]. However, maternal fasting hyperglycemia only explains 2-13% of variation in birth weight [10,11], and most LGA babies are not born to diabetic mothers [12]. This indicates that other factors are also important, for example, many LGA babies may be constitutionally large.

The contribution of common genetic variation to either SGA or LGA is not known, though associations between LGA and fetal genetic scores for birth weight have been demonstrated [13,14]. It is possible that a mismatch between a genetic score for birth weight and the actual, measured birth weight could help to identify babies who have either fallen short of, or exceeded, their growth potential. In clinical practice, this could be helpful if it improved the identification of FGR babies among those classified as SGA. Genetic studies of adult height previously investigated a similar question and showed that a genetic score composed of common variants was associated with adult height at the extremes of the population distribution [15]. However, the genetic score was not as extreme as expected in people with very short stature, suggesting that additional factors (e.g. rare mutations) were more important than common genetic variation in determining height in that group.

In the current study, we investigated the genetic contribution to SGA and LGA in mothers and babies of European ancestry, using common genetic variants identified in the most recent genome wide association study (GWAS) of birth weight [16]. The genetic variants at the 190 most strongly-associated loci have individually small effects, but collectively explain 7% of the variation in birth weight. These genetic variants have either a direct fetal effect on birth weight (i.e. those inherited by the fetus and acting via fetal pathways), or an indirect maternal effect (i.e. through a primary effect on the intrauterine environment), or some combination of the two (Fig. 1). The correlation between maternal and fetal genotypes means that associations between maternal genotype and birth weight can be confounded by fetal genotype effects, and vice-versa. To overcome this limitation, Warrington et al [16] estimated the independent maternal and fetal effect sizes at each of these loci. We used these independent effect sizes to calculate maternal and fetal genetic scores (GS) for birth weight. These GSs are designed to capture the independent maternal and fetal genetic contributions to birth weight. We tested for associations of these GSs with SGA and LGA. We then used simulations to test whether the GS in the SGA and LGA groups was consistent with the polygenic model. To investigate further the maternal genetic contribution to intrauterine factors known to be important across the birth weight distribution [16,17], we calculated genetic scores for fasting glucose (FG) and systolic blood pressure (SBP) and additionally tested their associations with SGA and LGA.

**Figure 1:**
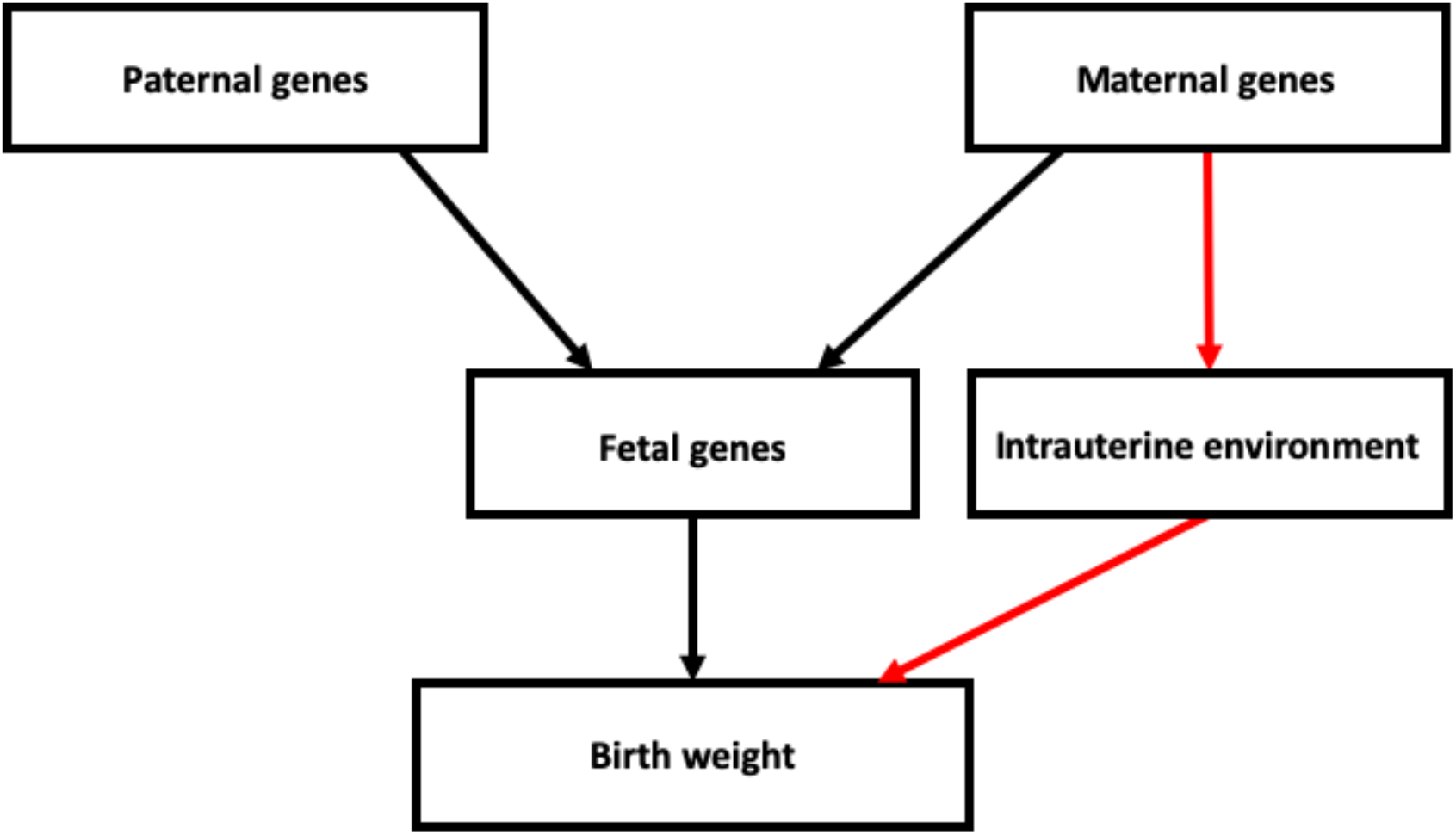
Diagram showing the possible pathways through which parental genotypes can influence birth weight. The black path represents direct fetal genetic effects on birth weight, and the red path represents maternal genetic factors which have an indirect effect on birth weight by modifying the intrauterine environment.

## Results

### Prevalence of SGA and LGA is strongly associated with maternal and fetal genetic scores for birth weight

The prevalence of SGA and LGA by percentile of fetal or maternal GS in ALSPAC (N=6,621) is shown in Figures S1 and S2, and the mean BW in ALSPAC by percentile of fetal and maternal GS is shown in Figure S3. Both the fetal and maternal genetic scores for birth weight showed strong associations with LGA and SGA (Figure 2, Table S4a) in the expected directions. A 1 decile increase in the fetal GS for birth weight was associated with a greater odds of LGA (OR=1.49 [95%CI: 1.40,1.58]; P=1.8×10^−42^) and a lower odds of SGA (OR=0.68 [0.64,0.72]; P=2.7×10^−36^). Similarly, the maternal GS showed strong associations with LGA (1.25 [1.17,1.33]; P=6.2×10^−12^) and SGA (0.80 [0.75,0.86]; P=4.6×10^−11^). These results were very similar to those of a sensitivity analysis of fetal GS conditional on maternal GS (and vice versa) in mother-child pairs (Table S4b), indicating that the use of the adjusted fetal and maternal weights [15] resulted in fetal and maternal GSs that were already independent of one another (Table S4).

**Figure 2:**
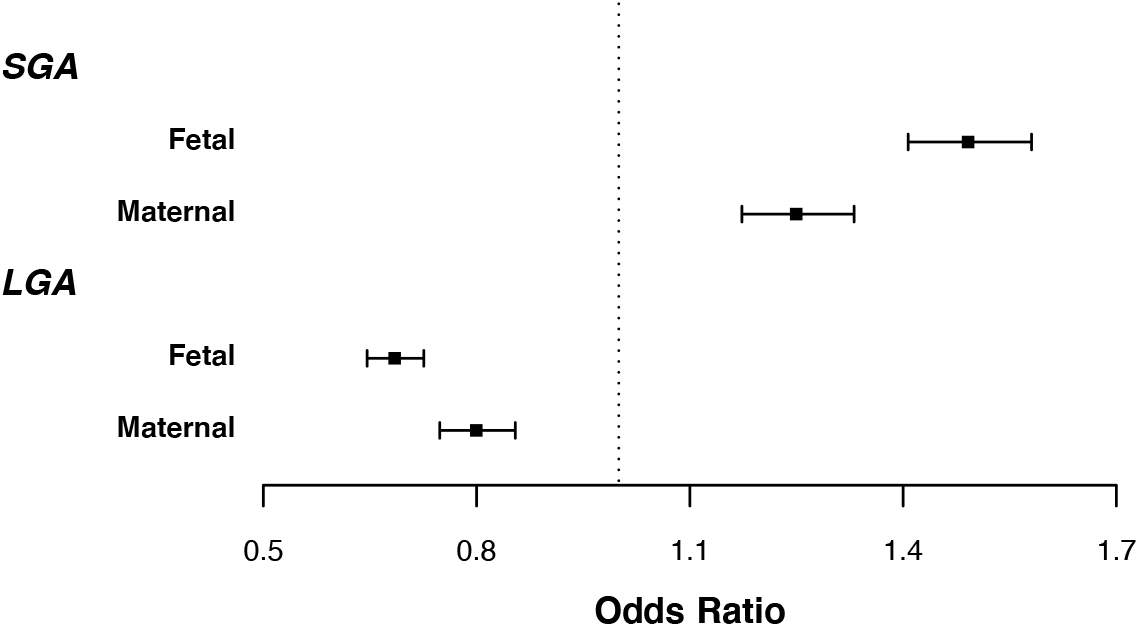
Odds of SGA or LGA per 1 decile higher fetal (N=12,125; ALSPAC, EFSOCH, NFBC66, NFBC86) or maternal (N=5,187; ALSPAC, EFSOCH) GS for birth weight. Error bars represent 95% confidence intervals, and weights used for fetal and maternal GS are independent of maternal and fetal effect respectively.

### Evidence that the GS in the lowest 3% of the population was higher than expected

Using simulation analyses in ALSPAC and EFSOCH (N=5,187) we identified evidence of deviation from the expected additive polygenic model in the lowest 10% bin of the phenotype distribution, which we then narrowed down to the lowest 3% bin for both the fetal and maternal GS for birth weight (P_fetal_=0.0034, P_maternal_=0.023; Figure 3; Table S5). The birth weight GS for both maternal and fetal genotype were higher than expected within this group indicating that there is a proportion of the SGA group whose birth weight is lower than would be expected given their GS.

**Figure 3:**
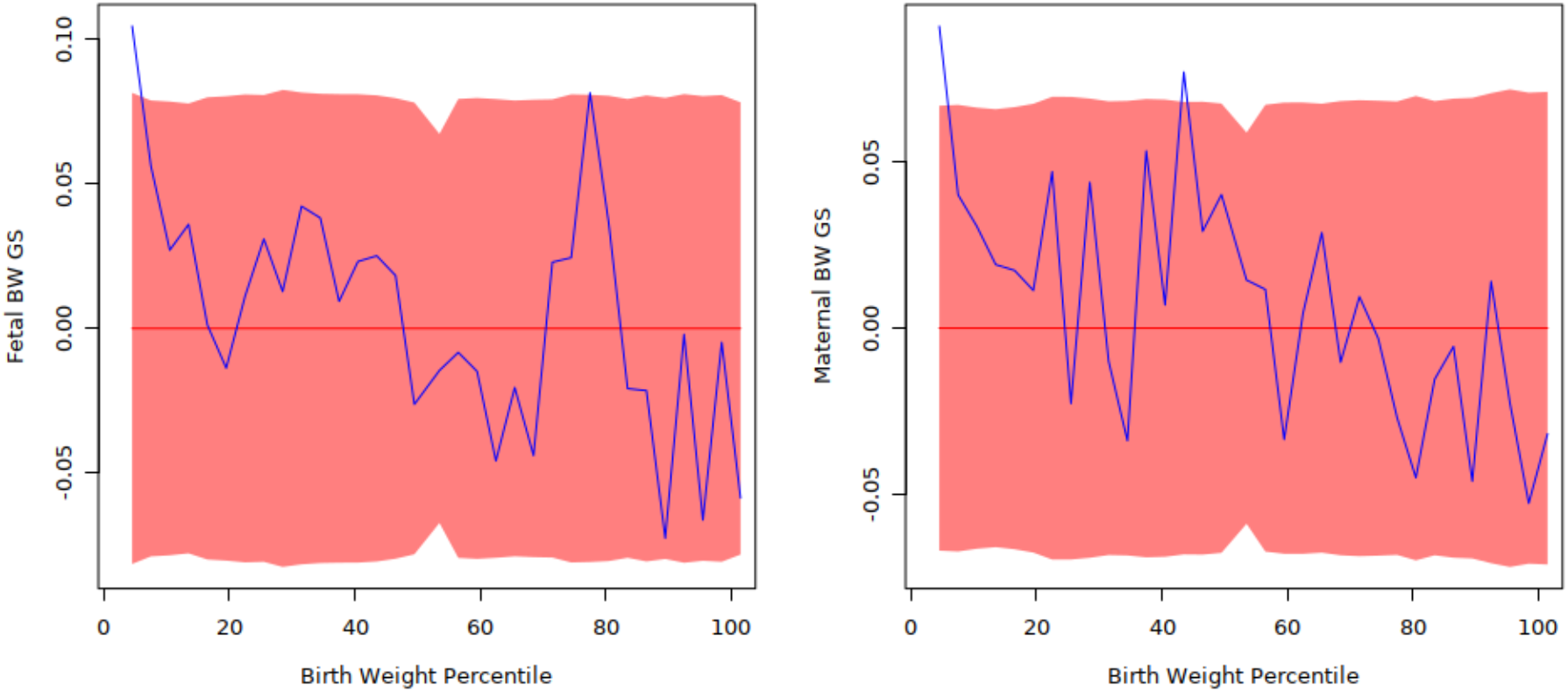
Difference between observed mean GS (blue line) and simulated mean of birth weight GS (under a fully polygenic model; red line) by 3% phenotype bins for fetal GS (left) and maternal GS (right) in ALSPAC. The shaded red area represents the 95% CI of the simulated mean.

There was no evidence of deviation from the expected additive polygenic model for the fetal GS in the top 10% of the phenotype distribution (P_fetal_= 0.1082). We saw some evidence of a lower maternal GS than expected in the top 10% group (P_maternal_=0.012) but in the top 3% of the phenotype distribution there was no evidence of deviation (P_maternal_=0.15; Table S5).

### Prevalence of SGA and LGA are associated with maternal genetic scores for fasting glucose and SBP

The maternal GS for fasting glucose was associated with higher odds of LGA (1.49 [1.19,1.87]; P=5.7×10^−4^), and the maternal SBP GS was associated with higher odds of SGA (1.31 [1.07,1.60]; P=9.5×10^−3^). These effects, observed across the genetic score distribution, are consistent with the known effects, respectively, of maternal gestational diabetes and hypertension on birth weight. Warrington et al [16] previously demonstrated continuous associations of both FG and SBP with BW in the healthy range. As a consequence of these continuous associations, we found some evidence of associations between lower genetic susceptibility to high FG and SBP and risk of SGA or LGA. Maternal fasting glucose GS was associated with lower odds of SGA (0.74 [0.58,0.94]; P=1.5×10^−2^). There was little evidence of an association between maternal SBP GS and LGA (0.87 [0.72,1.05]; P=1.4×10^−1^), although confidence intervals were wide and it would be informative to look at this association in studies with larger sample size (Figure 4, Table S6).

**Figure 4:**
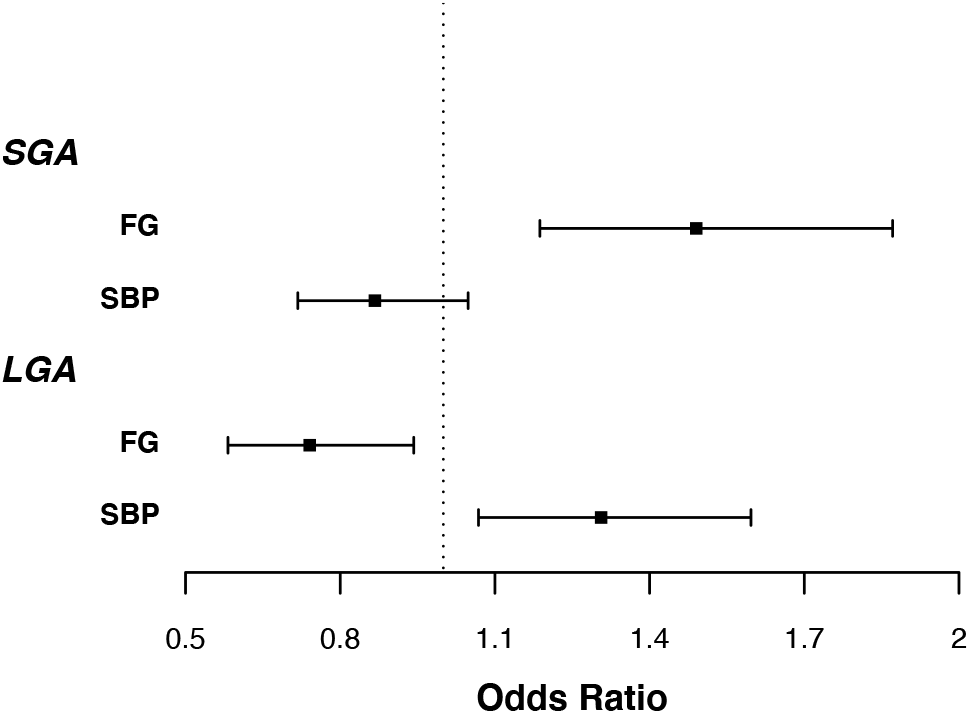
Odds of LGA/SGA 1 per decile higher maternal fasting glucose or SBP GS, corrected for fetal GS in ALSPAC and EFSOCH (N=5,187). Error bars represent 95% confidence intervals.

## Discussion

We have shown that common birth weight-associated genetic variation in both the mother and the fetus contribute to the probability that term infants will be born small or large for gestational age. While these results indicate that a large proportion of those infants classified as SGA/LGA represent the tail ends of the normal distribution of birth weight, we also found some evidence of an excess of individuals with a higher GS for birth weight than expected under the polygenic model in the bottom 3% of the birth weight distribution.

The terms IUGR and SGA are often used interchangeably [23], which can imply that the majority of SGA babies have experienced restriction to their uterine growth, although it has been estimated [6] that 50-70% of babies classified as SGA are constitutionally small. The strong association in our study between SGA and the fetal GS for birth weight, together with the deviation from the polygenic model in only the lowest 3% of babies, is consistent with any subset of growth-restricted babies being much smaller than those who are constitutionally small. Our results are also consistent with the observation that the risk of adverse outcomes increases with decreasing birth weight within the SGA group, and the highest rates of adverse outcomes are in those with birth weight below the 3^rd^ centile [24]. The higher than expected GS in the bottom 3% observed in the present study indicates that this group may be enriched for growth-restricted babies, although this finding would benefit from replication in larger studies.

We found no evidence that the fetal GS for birth weight deviated from the polygenic model in the top 10% of the birth weight distribution, but there was some evidence of deviation for the maternal GS in this group. When we looked in smaller percentile groups, however, there was no evidence of association in the top 3% of the distribution. Overall our results do not suggest strong deviation from the polygenic model in babies with high birth weights, but we cannot rule out that smaller deviations would be detectable in a larger sample.

Although we have used a population based definition of SGA, customised birth weight standards including maternal height, weight and ethnic group, for defining SGA have been shown to increase association between SGA classification and neonatal morbidity and perinatal death compared to using population based definitions [2], although it has been suggested that this could be an artefact due to inclusion of more preterm births classified as SGA under this definition [25]. Babies born preterm have increased risk of neonatal morbidity and mortality compared to term babies, and it is likely that different mechanisms affect growth in preterm babies compared to IUGR babies born at term. Additionally, individuals classified as SGA by population-based growth standards but not by customised standards are not at increased risk of perinatal mortality and morbidity compared to those of appropriate weight [26–28]. Including genetics in the definitions of SGA has the potential to further refine the identification of babies who have failed to properly reach their growth potential, as shown by the association between SGA and birth weight GS demonstrated here.

Fetal genetic effects on birth weight represent the constitutional growth potential of the fetus, while maternal genetics influence birth weight indirectly by modifying the intrauterine environment. The strong effects of the maternal GS for birth weight observed in our study indicate that maternal genetic variation acting via the in utero environment contributes to variation in SGA and LGA risk independently of the fetal genotype. Previous studies have shown that maternal FG and SBP have causal effects on birth weight in the normal range [15,16]. In women at highest genetic risk for raised FG and SBP, these could potentially contribute to LGA and SGA, respectively. We therefore investigated the associations between maternal genetic scores for FG or SBP and the risk of SGA or LGA. In line with known consequences of maternal gestational diabetes or hypertension, respectively, a higher maternal FG GS is associated with higher odds of LGA, while a higher maternal SBP GS increased odds of SGA. A lower GS for maternal FG was also associated with higher odds of SGA. This is consistent with naturally low maternal glucose levels contributing to SGA risk. A lower GS for maternal SBP showed weak evidence of association with LGA. Replication in larger studies will be necessary to determine the potential contribution of lower maternal blood pressure to LGA risk.

The current study has a number of strengths and limitations. Strengths include the fact that we have been able to construct independent maternal and fetal GS for birth weight. Separating maternal and fetal genetic contributions to birth weight is important because maternal and fetal genotype are correlated. This means that to avoid confounding, it is necessary to account for this correlation and obtain independent estimates of maternal and fetal genetic effects. Both a limitation and strength of our study is that all of the cohorts used were of European ancestry. There is a need for genetic studies in populations of ancestries other than Europeans, however, the GS were discovered in European ancestry individuals and are not necessarily applicable to other ancestry populations. A further limitation is that the cohorts analysed in this study contributed to the GWAS meta-analysis that identified SNPs contributing to the GS. However, these studies only constitute 3.8% of the fetal and 4.2% of the maternal samples included in the GWASs so any impact of winner’s curse on the associations in the current study is likely to be small. The use of only singleton, term babies in our analysis means that our results do not necessarily translate to pre-term babies or multiple births. While we had sufficient sample size to estimate the association of birth weight GS with SGA/LGA, the limited number of mother-child pairs which were available to examine the associations of FG and SBP GS with SGA/LGA meant that these estimates had large confidence intervals. Larger numbers of mother-child pairs would allow for more precise estimates, for example of the association of low SBP on LGA which has potentially clinically relevant implications. Although SGA and FGR are often used synonymously, there are differences between the terms. In our study do not have information required to distinguish FGR babies from SGA ones, meaning that we were not able to examine the association between BW GS and FGR specifically.

Our analysis has shown that common birth weight-associated genetic variation contributes to the risk of babies being born small or large for gestational age, and that the genetics underlying maternal fasting glucose and systolic blood pressure in the normal range also contribute to this risk. We found evidence of deviation from the polygenic model in the smallest 3% of babies, consistent with enrichment for fetal growth restriction in this group.

## Methods

### Cohort descriptions

Our analysis included a total of 6,938 individuals with birth weight and fetal genotype data, from 2 studies, plus 5,187 mother-offspring pairs with maternal and fetal genotype data and birth weight from 2 further studies. Studies are described below, and summary data is shown in Table 1.

**Table 1:** Descriptive statistics of studies contributing to analysis. EFSOCH and ALSPAC sample size is the number of mother-child pairs in the analysis. NFBC66 and NFBC86 sample size is the number of offspring in the analysis.

| Study | Country of origin | Year(s) of birth | Sample size (Male/Female (offspring sex)) | Data collection | Mean (SD) birth weight (grams) |  |  | Mean Maternal age (SD) | Mean Maternal BMI (SD) | Median (IQR) GA (weeks) at delivery |
| --- | --- | --- | --- | --- | --- | --- | --- | --- | --- | --- |
|  |  |  |  |  | Males | Females | Combined |  |  |  |
| ALSPAC | UK | ~1992 | 4570 (2262/3207) | Identified from obstetric data, records from the ALSPAC measurements, and birth notification | 3553 (491) | 3423 (450) | 3490 (476) | 28.0 (4.96) | 22.9 (3.83) | 40 (40-41) |
| EFSOCH | UK | 2000-2004 | 617 (322/294) | Measured within 12 hours of birth | 3585 (463) | 3447 (475) | 3519 (474) | 30.4 (5.28) | 28.0 (4.44) | 40 (39-41) |
| NFBC66 | Finland | 1966 | 3690 (1838/1851) | Measured in hospitals | 3607 (506) | 3480 (466) | 3541 (489) |  |  | 40 (39-41) |
| NFBC86 | Finland | 1986 | 3077 (1487/1589) | Measured in hospitals | 3626 (543) | 3519 (521) | 3572 (535) |  |  | 40 (39-40) |

#### ALSPAC

The ALSPAC (Avon Longitudinal Study of Parents and Children) is a longitudinal cohort study covering the area of the former county of Avon, UK [18,19]. Women who were pregnant, living in the study area and had an expected delivery date between 1 April 1991 and 31 December 1992 were eligible for inclusion in the study. Birth weight and related data were abstracted from medical records. Children were genotyped on the Illumina HumanHap550 chip and mothers were genotyped using the Illumina human660W quad chip. QC was undertaken as described previously [16] and imputation was to the HRC reference panel yielding data available for 8884 mothers and 8860 children with genotype data available. Of these, 4570 mother-child pairs with phenotype data were available for analysis.

#### EFSOCH

The Exeter Family Study of Childhood Health (EFSOCH) is a prospective study of children born between 2000 and 2004 in the Exeter region, UK, and their parents [20]. Birth weight measurements were performed as soon as possible after delivery. Individuals were genotyped using the Illumina Infinium HumanCoreExome-24 array. Genotype QC and imputation have been described previously [13]. After genotype QC, 938 mothers and 712 children with genotype data remained. Of these, 617 mother-child pairs with phenotype data were available for analysis.

#### NFBC66

The Northern Finland Birth Cohort 1966 (NFBC66) consists of mothers expected to give birth in the provinces of Oulu and Lapland in 1966 [21]. Birth weight and related characteristics were measured by examinations occurring directly after birth. Samples were genotyped on the Infinium 370cnvDuo. QC and imputation to the 1000 genomes reference panel were done centrally, resulting in 5402 children with genotype data. A total of 3690 children with phenotype data were available for analysis.

#### NFBC86

The Northern Finland Birth Cohort 1986 (NFBC86) consists of individuals born in the northern-most provinces of Finland between 1^st^ July 1985 and 30^th^ June 1986. Birth weight data was collected from hospital records after delivery. Following genotype QC, 3742 children with genotype data were available, with 3248 of these having phenotype data [22].

### Phenotype definitions

Since different mechanisms may lead to SGA and LGA in term and preterm infants, and in multiple births, we focused on term, singleton infants. Within each cohort, birth weight was regressed against sex and gestational age in term births (gestational age >= 37 weeks), and residuals from the regression model were calculated. Individuals with the lowest and highest 10% of this adjusted birth weight within each cohort respectively were defined as SGA and LGA. Controls for comparison with SGA were taken as birth weight >=10%, and for comparison with LGA as birth weight <=90%.

Genetic scores (GS) for birth weight, fasting glucose (FG) and systolic blood pressure (SBP) were calculated for all included individuals in each cohort. GS were calculated using equation (1) where N_SNP_ is the total number of SNPs, w_i_ is the weight for SNP i and g_i_ is the genotype at SNPi. The same SNPs were included in the maternal and fetal GS. The SNP weightings, w_i_, were taken from the structural equation model-adjusted effect estimates in [16]. These adjusted effect estimates are designed to capture the independent maternal and fetal contribution at each SNP, which would otherwise be confounded by the correlation between maternal and fetal genotype. For the weights in the FG and SBP scores, we used effect estimates reported in large GWAS of FG and SBP, respectively, and the same weights were used for both maternal and fetal GS. SNPs and weights used in each score are given in Tables S1-3.

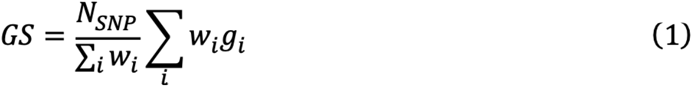

### Association analysis

#### Associations between SGA/LGA and maternal or fetal genetic scores for birth weight

Fetal genotypes (total n=12,125) were available in ALSPAC (N=4,570), EFSOCH (N=617), NFBC66 (N=3,690) and NFBC86 (N=3,248). Associations were tested between the fetal GS for birth weight and outcomes (SGA/LGA) using logistic regression, including sex and gestational age as covariates to control for residual confounding. Maternal genotypes were also available in ALSPAC (N=4,570) and EFSOCH (N=617). In these cohorts, associations between SGA/LGA and maternal GSs for birth weight were also tested in analogous regression models. Additional analyses including both maternal and fetal GS in the same regression model were performed to control for any residual correlation between maternal and fetal genotype. Results were meta-analysed using inverse variance weighted meta-analysis.

#### Investigating Deviations from the expected Polygenic Model in SGA/LGA

We performed simulations to assess whether there was any evidence of deviation from the expected polygenic model. Briefly, within ALSPAC and EFSOCH we calculated the associations of each fetal and maternal SNP with birth weight within each cohort, adjusted for maternal and fetal genotype, respectively. We simulated genotypes under the allele frequencies observed within each cohort. GSs were calculated for each simulated individual using simulated genotypes weighted by the within-cohort effect sizes. Phenotypes were then simulated as

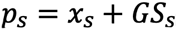

where p_s_ is the simulated phenotype for individual s, GS_s_ is the simulated genetic score for sample s and x_s_ is sampled from a normal distribution

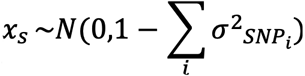

where 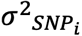 is the variance explained by SNP i. The mean GS within each bin of simulated phenotype was calculated. Simulations were performed 10,000 times to generate a simulated expected distribution of GS under an additive polygenic model. The observed mean GS was then compared to the expected distribution and an empirical p value was calculated. These p values were then meta-analysed.

#### Associations between SGA/LGA and maternal genetic scores for fasting glucose or systolic blood pressure

In ALSPAC and EFSOCH, where maternal genotypes were available, we tested the associations between the maternal GSs for FG and SBP, with outcomes (SGA/LGA) using linear regression, including sex and gestational age as covariates to control for residual confounding. Additional analyses including both maternal and fetal GS in the same regression model were performed to control for the correlation between maternal and fetal genotype. Results were meta-analysed using inverse variance weighted meta-analysis.

## Acknowledgements

**ALSPAC** Ethical approval for the study was obtained from the ALSPAC Ethics and Law Committee and the Local Research Ethics Committees.

We are extremely grateful to all the families who took part in this study, the midwives for their help in recruiting them, and the whole ALSPAC team, which includes interviewers, computer and laboratory technicians, clerical workers, research scientists, volunteers, managers, receptionists and nurses.

The UK Medical Research Council and Wellcome (Grant ref: 217065/Z/19/Z) and the University of Bristol provide core support for ALSPAC. This publication is the work of the authors and Rachel M. Freathy will serve as guarantor for the contents of this paper.

Please note that the study website contains details of all the data that is available through a fully searchable data dictionary and variable search tool: http://www.bristol.ac.uk/alspac/researchers/our-data/.

**EFSOCH**: The Exeter Family Study of Childhood Health (EFSOCH) was supported by South West NHS Research and Development, Exeter NHS Research and Development, the Darlington Trust and the Peninsula National Institute of Health Research (NIHR) Clinical Research Facility at the University of Exeter. The views expressed are those of the authors and not necessarily those of NIHR, the NHS or the Department of Health. Genotyping of the EFSOCH study samples was funded by the Welcome Trust and Royal Society grant WT104150.

**NFBC 1966 and 1986**: We thank the late professor Paula Rantakallio (launch of NFBC1966), the participants in the 31- and 46-years-old study and the NFBC project center. We thank professor Anna-Liisa Hartikainen (launch of NFBC1986), the participants in the study and the NFBC project center.

The Northern Finland Birth Cohort (NFBC) 1966 and 1986 studies have received financial support from the Academy of Finland (project grants 104781, 120315, 129269, 1114194, 24300796, Center of Excellence in Complex Disease Genetics and SALVE), University Hospital Oulu, Biocenter, University of Oulu, Finland (75617), NIHM (MH063706, Smalley and Jarvelin), Juselius Foundation, NHLBI grant 5R01HL087679-02 through the STAMPEED program (1RL1MH083268-01), NIH/NIMH (5R01MH63706:02), the European Commission (EURO-BLCS, Framework 5 award QLG1-CT-2000-01643), ENGAGE project and grant agreement HEALTH-F4-2007-201413, EU FP7 EurHEALTHAgeing-277849, the Medical Research Council, UK (G0500539, G0600705, G1002319, PrevMetSyn/SALVE) and the MRC, Centenary Early Career Award. The program is currently being funded by the H2020 DynaHEALTH action (grant agreement 633595) and academy of Finland EGEA-project (285547). The DNA extractions, sample quality controls, biobank up-keeping and aliquotting was performed in the National Public Health Institute, Biomedicum Helsinki, Finland and supported financially by the Academy of Finland and Biocentrum Helsinki. We thank the late Professor Paula Rantakallio (launch of NFBCs), and Ms Outi Tornwall and Ms Minttu Jussila (DNA biobanking). The authors would like to acknowledge the contribution of the late Academian of Science Leena Peltonen.

---

**Figure S1:**
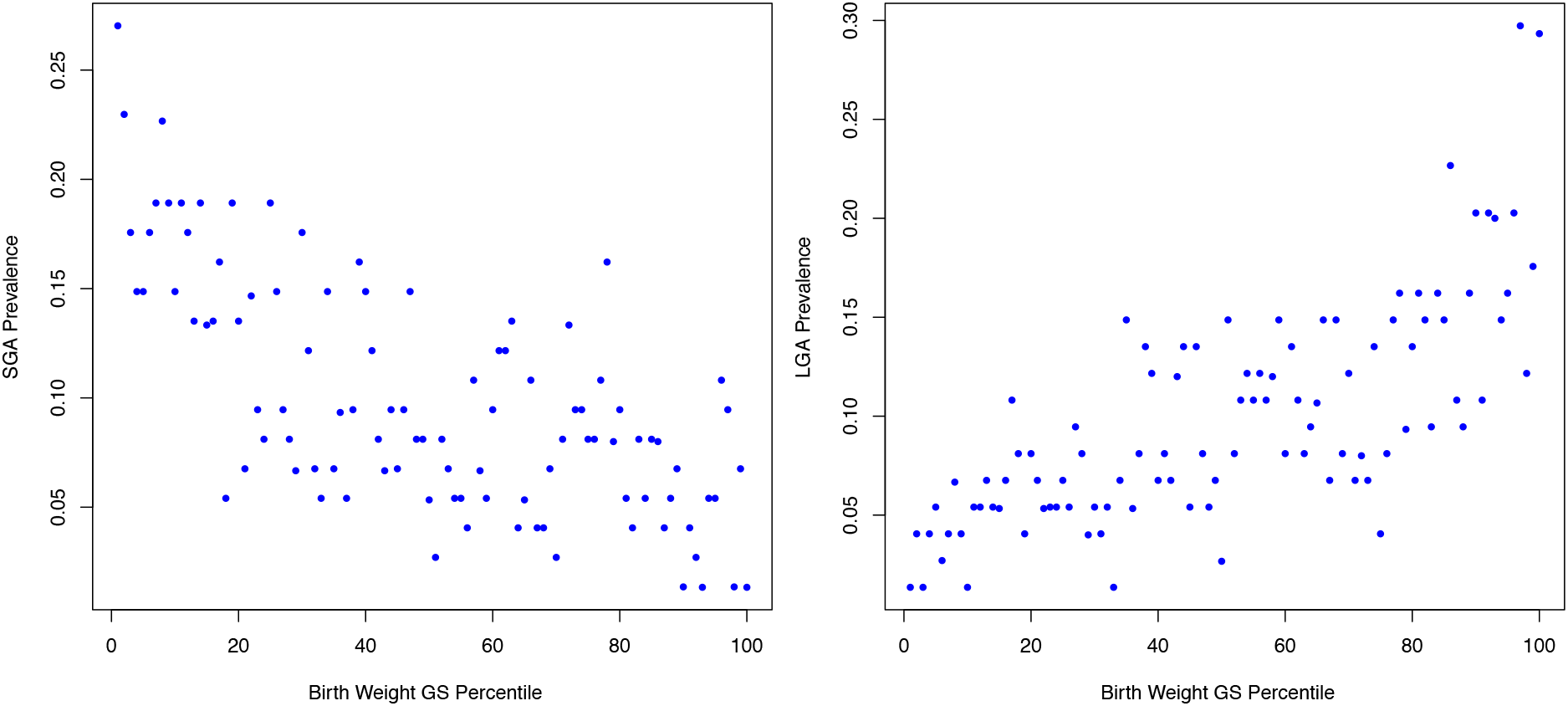
Fraction of babies born SGA (left) or LGA (right) by percentile bins of fetal birth weight GS in ALSPAC (N=6,621)

**Figure S2:**
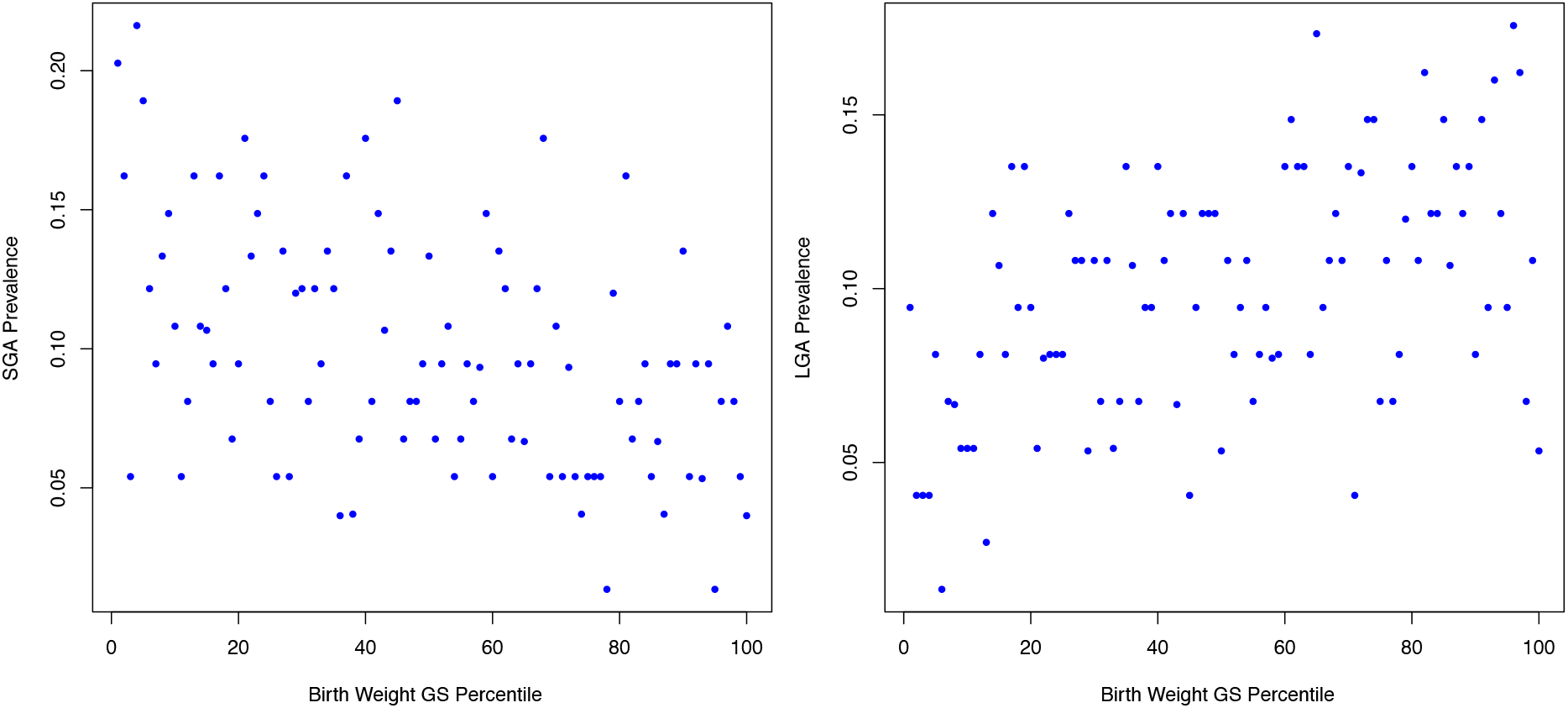
Fraction of babies born SGA (left) or LGA (right) by percentile bins of maternal birth weight GS in ALSPAC (N=6,621)

**Figure S3:**
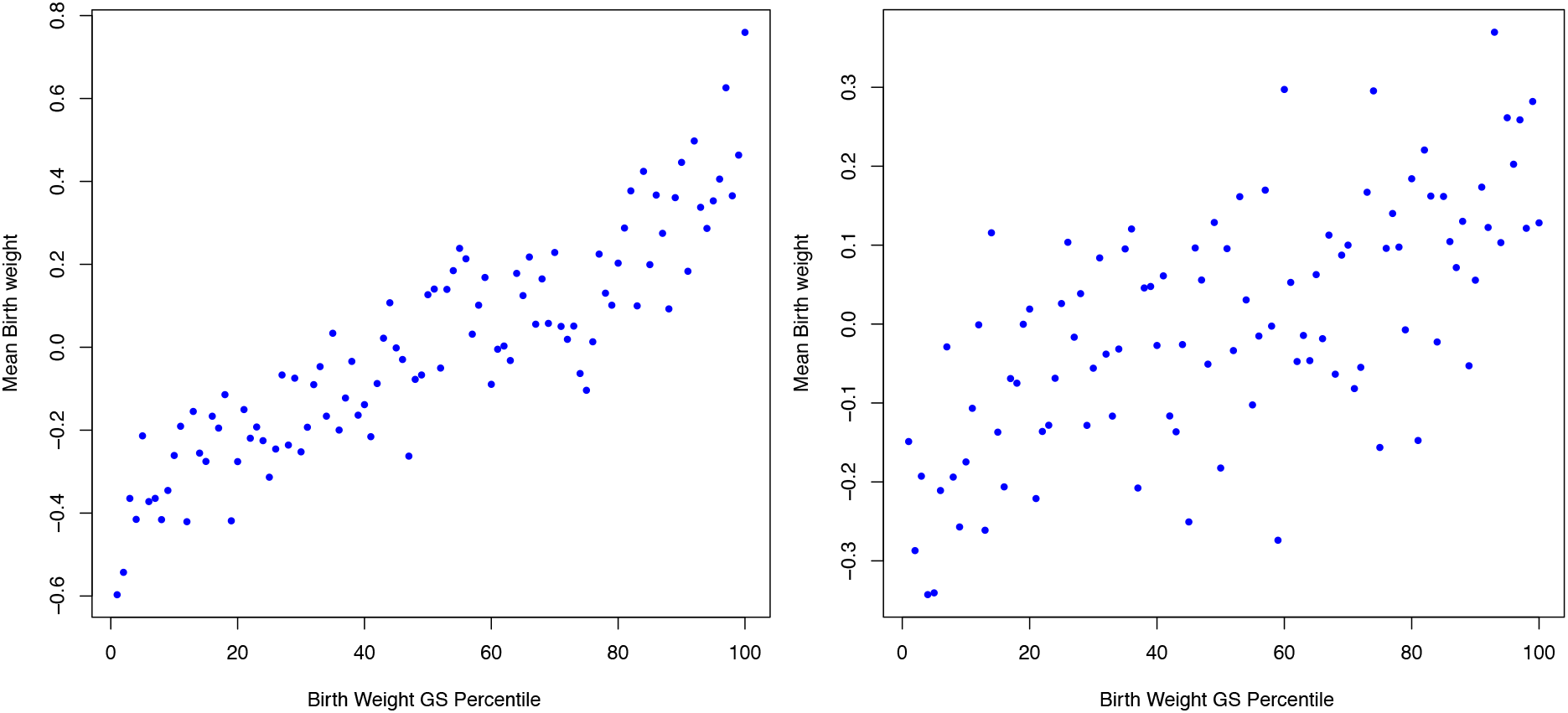
Mean birth weight in ALSPAC by percentile bins of fetal (left) and maternal (right) birth weight GS (N=6,621)

